# Fast near-whole brain imaging in adult Drosophila during responses to stimuli and behavior

**DOI:** 10.1101/033803

**Authors:** Sophie Aimon, Takeo Katsuki, Tongqiu Jia, Logan Grosenick, Michael Broxton, Karl Deisseroth, Terrence J. Sejnowski, Ralph J. Greenspan

## Abstract

Whole brain recordings give us a global perspective of the brain in action. In this study, we describe a method using light field microscopy to record near-whole brain calcium and voltage activity, at high speed, in behaving adult flies. We first obtained global activity maps for various stimuli and behaviors. Notably, we found that brain activity increased on a global scale when the fly walked but not when it groomed. This global increase with walking was particularly strong in dopamine neurons. Second, we extracted maps of spatially distinct sources of activity as well as their time series using principal component analysis and independent component analysis. The characteristic shapes in the maps matched the anatomy of sub-neuropil regions and in some cases a specific neuron type. Brain structures that responded to light and odor were consistent with previous reports, confirming the new technique’s validity. We also observed previously uncharacterized behavior-related activity, as well as patterns of spontaneous voltage activity.

## Introduction

Measuring activity at the scale of the whole brain is critical to understanding how different brain regions interact to process and control sensory inputs, internal states, and behavior. Whole-brain recordings not only reveal which regions are involved in which functions and with what dynamics, but also help interpret the effects of a targeted intervention (e.g. a lesion or local alteration with optogenetics) on the whole network, and give context to local electrophysiology recordings. However, techniques for imaging a whole brain so far have been orders of magnitude slower than neuronal electrical activity. In fact, recent reports of volumetric whole brain fluorescence imaging in Zebrafish and Drosophila larvae had a frame rate of 12 Hz [1] and 5 Hz [2], respectively. By constrast, light field microscopy [3–9], makes it possible to image large volumes of scattering brain tissue at more than 100 Hz.

In this study, we leverage this technique to record large-scale activity in the brain of behaving adult fruit flies. We present a method to optically access the fly’s brain while enabling it to retain the ability to walk or groom. We show that the near-whole brain can be imaged with a 20x objective at a frame rate up to 200 Hz and fluorescence recorded from pan-neuronally expressed calcium (GCaMP6 [10]) or voltage (ArcLight [11]) probes. We present rich datasets of near-whole brain activity and behavior, as well as two analysis methods. First, we map activity for specific stimuli and behaviors with short time-scales; for example, we compared activity when the fly rested, walked, and groomed. Second, we apply a computational method (principal component analysis, or PCA, followed by independent component analysis, or ICA) to extract components representing spatially distinct sources of activity [6,12,13]. We show that these sources correspond to sub-neuropil areas or processes from small populations of neurons that are anatomically well characterized, and we compare their responses to flashes of light or odor puffs with those in literature reports of experiments done on restricted regions. Additionally, by using this method, we discovered neuronal projections whose activity correlated with turning, as well as previously unreported patterns of spontaneous voltage activity.

## Results

### Imaging the near-whole brain of behaving adult Drosophila

We first fixed a fly’s head by rotating it 45 degrees or more around the transversal axis to decrease the depth of the volume imaged and to improve access to the brain. We then exposed the brain while keeping the eyes, antennae, and legs intact and clean (see methods section and Fig. S1). A ball was placed under the fly’s tarsi so that it could typically rest, walk, and groom. We imaged the fly brain’s fluorescence using light field microscopy. As shown in Fig 1A, we modified an upright epifluorescence microscope (equipped with a 20x 1.0 NA or a 40 x 0.8 NA objective) by adding a microlens array at the image plane of the objective and placing the camera sensor at the image plane of the microlens array through relay lenses. We recorded light field images continuously with a high-speed sCMOS camera up to 100 Hz for GCaMP6 and 200 Hz for ArcLight (using the middle of the camera sensor).

**Figure 1:**
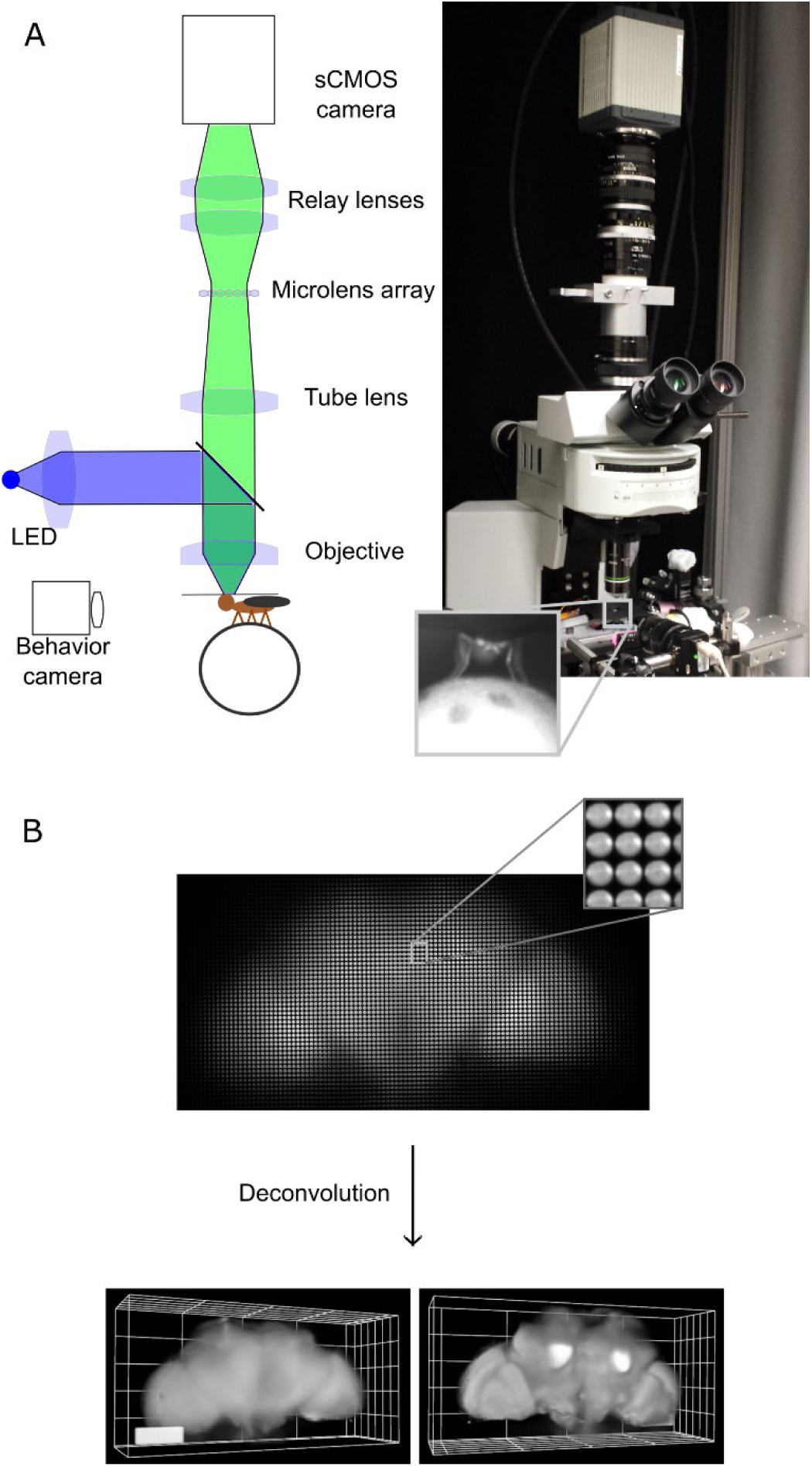
Imaging the brain of adult behaving flies using light field microscopy. A) Experimental set-up. The fly is head-fixed and its tarsi are touching a ball. The light from the brain’s fluorescence goes through the objective, the microscope tube lens, a microlens array, and relay lenses, onto the sensor of a high-speed sCMOS camera. Another camera in front of the fly records its behavior. B) Example of a light field deconvolution (fly’s genotype: nsyb-Gal4, UAS-ArcLight). Top: 2D light field image acquired in 5ms—one camera acquisition period—with a 20x NA 1.0 objective. Bottom: Anterior and posterior views (slightly tilted sideways) of the computationally reconstructed volume. 3D bar is 90×30×30 microns. See also Figs. S1, S2, and S3.

We then reconstructed the volumes—typically 600 × 300 × 200 µm^3^ to encompass the whole brain (Fig. 1B)—using the volumetric deconvolution method for light field microscopy described in ref. [3]. Note that unlike other microscopy techniques that are based on scanning (e.g. two-photon, confocal or light sheet microscopy), excitation light illuminates the entire brain all the time, and all the photons emitted in the numerical aperture of the objective are used to reconstruct the image (minus a ∼40% loss through the objective, tube lens, micro-lens array and relay lenses). This maximizes the number of photons collected (and thus information about brain activity) per unit of volume and time.

We used 2 µm fluorescent beads embedded in a gel to measure the point spread function, and found that it widens with distance from the focal plane, varying from 3.5 to 12 µm laterally and from 6 to 35 µm axially for 20x 1.0 NA objective, and varying from 2 to 7 µm laterally and from 4 to 22 µm axially for 40x 0.8 NA objective (Fig. S2 and theoretical expression in [3]). As shown in Fig. S3 (presenting ArcLight’s baseline fluorescence) and below, this resolution was sufficient to recognize neuropil structures and extract activity from sub-neuropil compartments.

### Global activity during response to stimuli and behavior

Movies 1 and 2, present maximum z projections of near-whole brain activity (after preprocessing as described in Fig. S4 and the Methods section), when stimuli were presented to the fly. Figs. 2C and Fig S5C show maps of the response to stimuli. We found strong increases in activity at the onset of puffs of odor and flashes of UV light in specific parts of the brain (see Figs. S5, S6, and S7), in accordance with previous reports in the literature: the strongest responses to light involved mostly the optic lobes, optical glomeruli, and posterior slope, and the responses to odor involved the antennal lobe and most of the dorsal neuropils including the lateral horn, superior neuropils and the mushroom body (Fig. S5C, S6 and S7). The global map of the response to stimuli was similar for calcium (GCaMP6) and voltage (ArcLight) activity (Fig. S5C).

**Figure 2:**
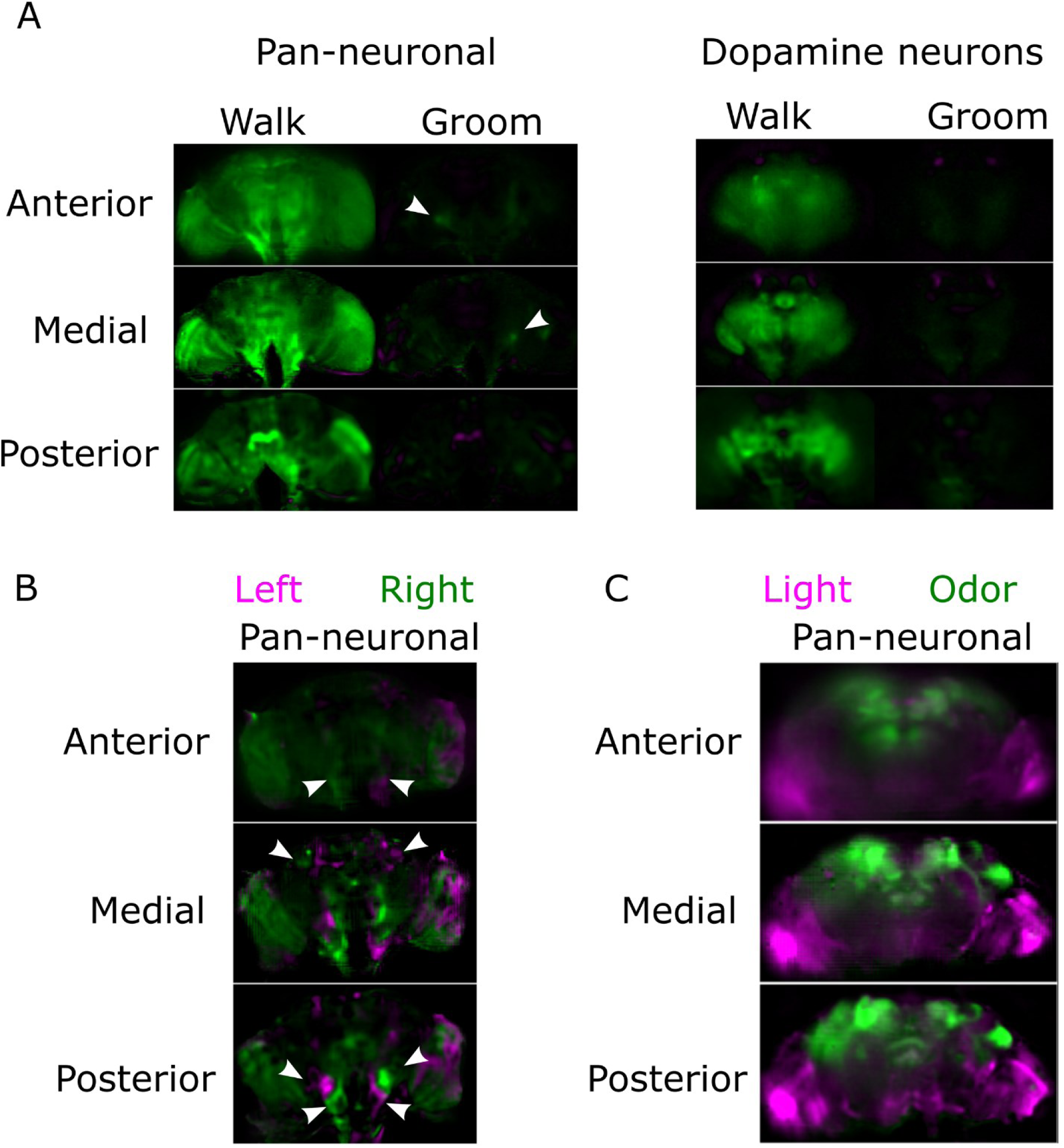
Near-whole brain activity maps for various conditions. A) Comparison of fluorescence intensity when the fly rests and when it is active (either walking or grooming). The pixel value is green if the fluorescence is higher during the behavior, and magenta if the fluorescence is higher during rest. The arrows point to regions active during grooming that were reproducible from fly to fly. Both flies expressed GCaMP6f. The pan-neuronal driver was nsyb-Gal4 and the dopamine driver was TH-Gal4. Note that there is an angle (39 degrees for the pan-neuronal and 20 degrees for the dopamine neurons) between the Z-axis and the anterior/posterior axis. B) Comparison of fluorescence intensity when the fly turns left or right. The arrows point to regions symmetrical and reproducible from fly to fly. The fly’s genotype was nsyb-Gal4 and UAS-GCaMP6f. Note that there is a 40 degrees angle between the Z-axis and the anterior-posterior axis. The pixel value is green if the fluorescence is higher during turning left, magenta if the fluorescence is higher during turning right. C) Comparison of fluorescence increase during the response to stimuli for odor (magenta), and light (green). The fly’s genotype was Cha-Gal4, GMR57C10-Gal4, and UAS-GCaMP6f. Note that the brain is tilted 19 degrees along the lateral axis. See also Figs. S4–8.

We also examined near-whole brain activity in the absence of external stimuli (i. e., spontaneous behavior), which consisted of walking, grooming, and resting (see Movies 3-7). Most strikingly, the brain was more active on a global scale when the fly walked than when it rested or groomed (see Movie 3, Figs. 2A, S5A, and S8). To verify that this response was linked to walking rather than the optic flow from the ball, we repeated the experiment with a blind *norpA* mutant fly, and again found a global increase during walking in comparison with rest (see Movie 4 and Fig. S5A). In contrast, we found only local activation in the region of the saddle, wedge and antennal mechanosensory and motor center (in 4 out of 5 flies) during grooming. To investigate whether this global increase was coming exclusively from neurons expressing one type of neurotransmitter or neuromodulator, we performed the same experiments with more restricted lines. We also found a global increase when GCaMP6f was expressed in cholinergic neurons (the majority of excitatory neurons in the fly brain) (Movie 5 and Fig S5A). When GCaMP6f was expressed in dopamine neurons only (Movie 6, Fig 2A and Fig S5A), we observed a strong large-scale increase of activity tightly locked with walking. Surprisingly little activity during resting or grooming, apart from the mushroom body compartments. We observed more activity unrelated to behavior in flies expressing GCaMP6f in both dopamine and serotonin neurons (Movie 7).

Figs. 2B and FigS5B show the difference in activity when the fly turned left compared to right. Although there was a strong variability from fly to fly that will need to be characterized in future studies, we observed antisymmetric patterns in the ventral areas and in the lateral superior protocerebrum (as indicated by the arrows) in all flies.

### Source extraction algorithm reveals functional maps at the sub-neuropil level

We used a combination of statistical methods to extract maps and time series of spatially distinct sources of activity (see Methods section for details). Briefly, we first applied PCA (using singular value decomposition) to find maps of correlated activity and to reduce dimensionality. We then applied ICA to the PCA maps to find sparse functional regions (see analysis pipeline in Fig. S4). Fig. 3 shows z-stacks containing all components (z-score > 3) from pan-neuronal GCaMP6 recordings (see also additional flies in Figs. S9 and S10). We also performed the same analysis with flies that pan-neuronally expressed activity-independent GFP to help identify artifact-related components—movement, aliaising, or noise as shown in Fig. S11.

**Fig. 3:**
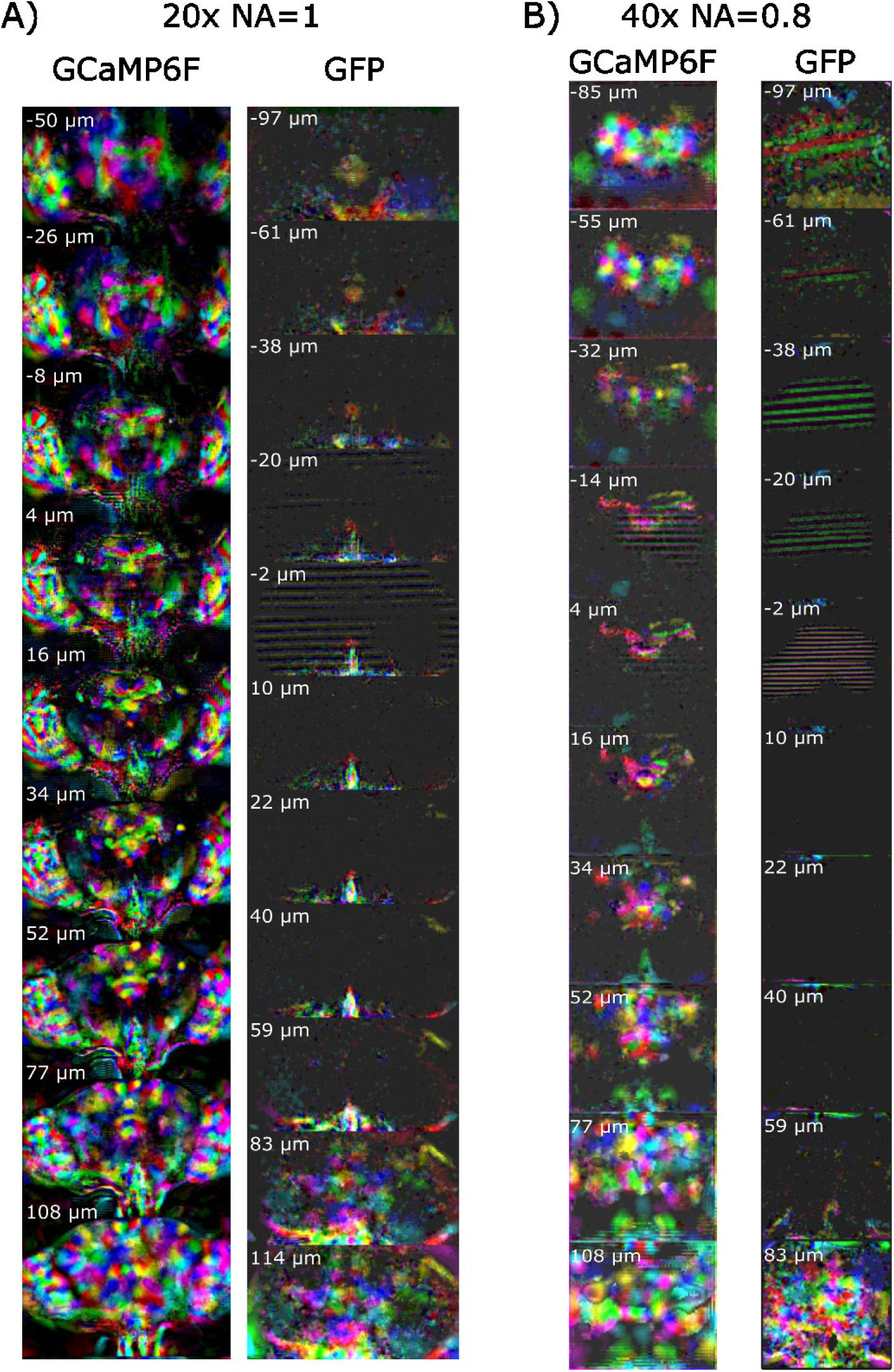
Z-stack slices of the 3D map for all the components extracted using PCA/ICA. The maps from calcium activity (GCaMP6f) are compared with the maps obtained with a control independent of activity (GFP). Different colors correspond to different components, which were assigned randomly. Note that the slice depth is larger when it is farther from the middle of the brain (see Methods section). See Figs. S9 and S10 for maps aligned to an anatomical template, and see S11 for examples of individual artefactual components.

Even though PCA and ICA are mathematical algorithms that make minimal assumptions about the brain, most functional maps matched well with the anatomical structures. We aligned the brain with an anatomical template [14] using landmarks registration to automatically sort the components by brain region (Fig. 4A). In Fig. 4B, the left column presents the component’s thresholded maps, whereas the right column presents central complex structures from (ref [15]) or neuronal processes from (ref. [16]) assembled using Virtual Fly Brain [17]. Several sub-neuropil regions are recognizable from the shape of the maps (e.g., protocerebral bridge glomeruli, ellipsoid body rings and fan-shaped body layers).

**Figure 4:**
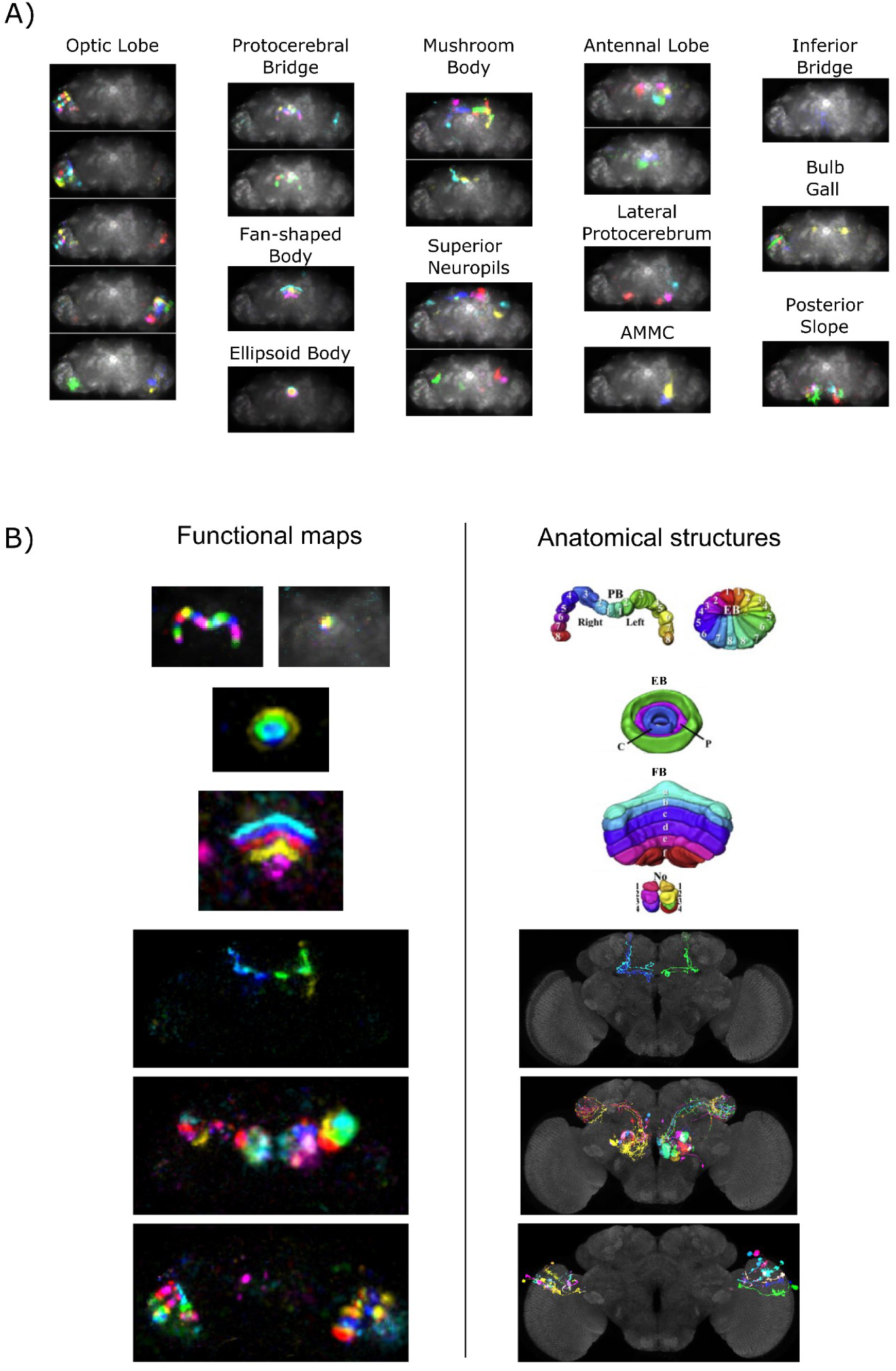
A) Components automatically sorted by regions containing the highest average in the map, and projected along the z-axis. Note that this sorting algorithm could be inaccurate in the case of maps containing small parts of large regions and noise in very small regions (i.e., the bulb or gall). B) Comparison between functional and anatomical maps. Left: functional maps from a pan-neuronal (GMR57C10-Gal4 and Cha-Gal4) GCaMP6f experiment. Right: corresponding anatomical structures. The top three images correspond to central complex structures–modified from Cell Reports, Vol. 3, Lin, C. Y. et al.,“ A Comprehensive Wiring Diagram of the Protocerebral Bridge for Visual Information Processing in the Drosophila Brain.,” 1739-1753, Copyright 2013, with permission from Elsevier, and the four bottom images were neurons from the Virtual Fly Brain database. The brain for the functional data is tilted 19 degrees along the lateral axis compared to the template presented on the right. Note that the functional maps in the fan-shaped body and the lateral horn are the same scale as functional maps obtained with higher resolution microscopy techniques ([36] and [37], respectively). See also Figs. S11 through S15 for considerations regarding the associated time series.

For some components, parts of a specific neuron type were identifiable from the combination of sub-neuropil regions present in the map. For example, z-scored maps containing signals in one antennal lobe glomerulus, in the calyx, and in the lateral horn, likely resulted from the activity of one type of antennal lobe projection neurons. Likewise, z-scored maps with signals spanning both the horizontal and vertical mushroom body lobes likely resulted from activity in alpha-beta or alpha’-beta’ Kenyon cell axons, whereas maps with signal in the horizontal lobe only likely resulted from activity in the gamma Kenyon cell axons. The maps of the components matching the protocerebral bridge glomeruli also often contained radial parts of the ellipsoid body (e.g., Fig.4B, left column top panel), suggesting that these components might originate from tile or wedge neurons [15,18].

### Extracted sources’ time series retrieve known physiology in response to stimuli, and reveal projections involved in turning left or right during walking

The component’s time series (resulting from PCA/ICA or from ROI ΔF/F averages: Fig. S12 and S14 to S16) were consistent with previous reports of activity from the brain structures identified in the components’ maps. Most of the components responding to the onset and/or offset of light were in the optic lobe [19] (Figs. S13 and S14). In contrast, components responding to puffs of odors were mostly in the antennal lobe, the lateral horn, and the mushroom body [20] (Figs. S12 and S15). Components likely representing the activity of antennal projection neurons were spontaneously active in the absence of odor, but their activity strongly increased with the odor (Fig S15 panel A), consistent with the literature [21].

In addition, regions in the lateral protocerebrum, the superior slope, the AMMC (antennal mechanosensory and motor center), the saddle, and the protocerebral bridge were strongly active when the fly walked (Fig. S16). This is consistent with previous anatomical studies; the projection from the descending neurons are most dense in the posterior slope, ventrolateral protocerebrum, and the AMMC [22]. Some of these walking-related components were strongly correlated with turning left or right (Fig. 5). We found the same components when using a *Cha-Gal4* driver instead of a pan-neuronal driver (see Fig. S17), which suggests that all those neurons are cholinergic. Their strong structural characteristics (e.g., small neuropil areas forming an inverted “V” shape, fine tracts) will help identify candidate drivers and neurons in anatomy databases in follow up studies. Note that these components are mostly present in the posterior slope, as are neurons involved in turning during flight [23].

**Figure 5:**
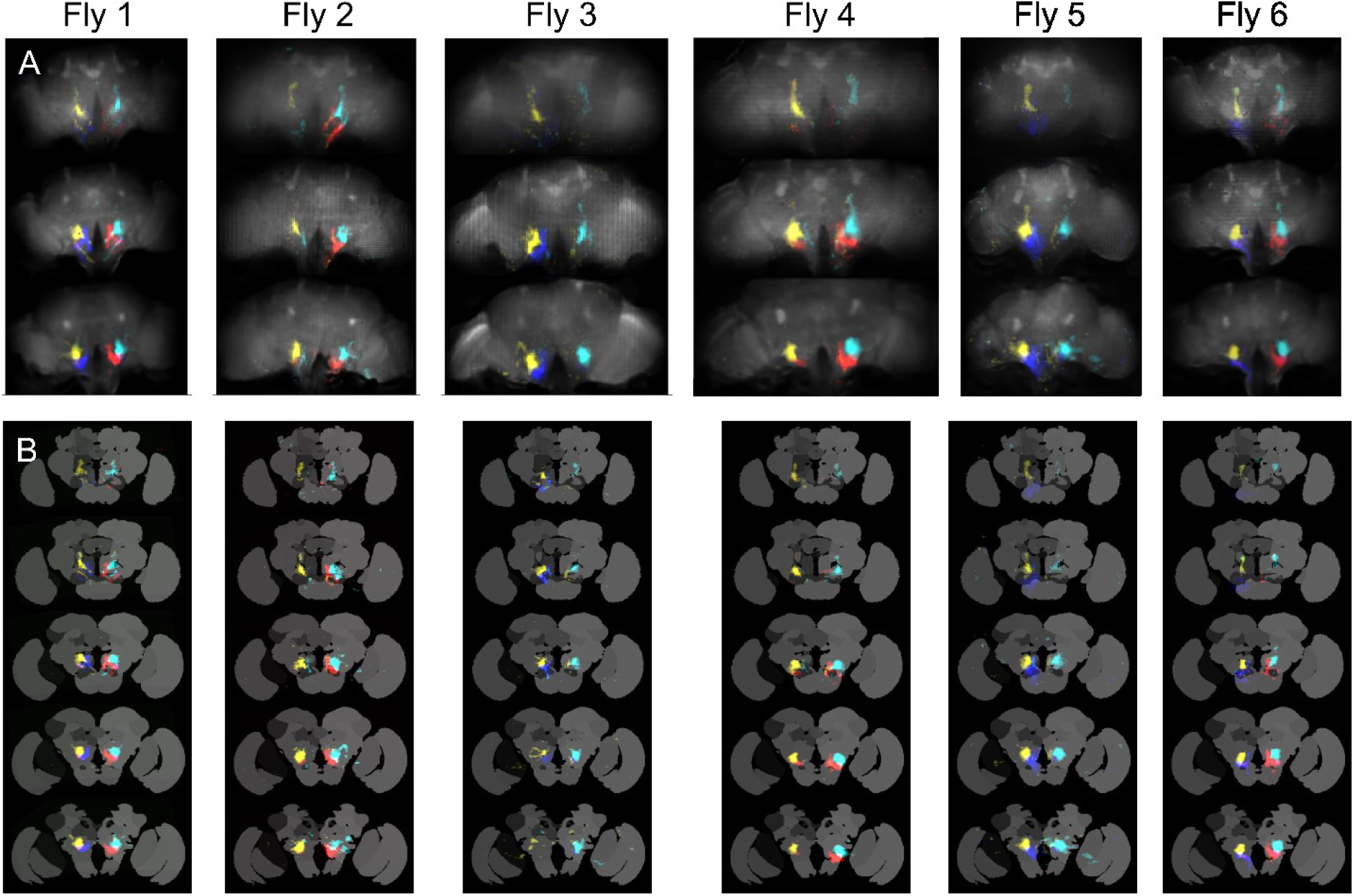
Z-stacks of components among the 3 to 6 most correlated to turning left (cyan and blue) or right (red and yellow). Each color corresponds to one component. Flies expressed GCaMP6 pan-neuronally. A) Anterior, medial and posterior views in the original configuration. B) The same components aligned to an anatomical template’s z-stack. See also Fig. S16.

### Restricted drivers’ functional maps match single neuron’s anatomy

Figs. 6 and S18 show components obtained when using more restrictive drivers for dopamine neurons: *TH-Gal4* (data are the same as in Movie 6). As before, we sorted the components’ traces and maps by brain regions (Fig. 6A). In agreement with our observation from Movie 6, most components were tightly correlated with the fly walking (forest green traces interleaved with the components’ traces). Fig. 6B reproduces some of the maps in Fig. 6A, along with anatomical maps of single dopaminergic neurons from the Virtual Fly Brain database [17]. Some maps had unequivocal anatomical counterparts. For example, the first two maps matched well with the anatomy of processes from dopaminergic PPL1 neurons innervating mushroom body alpha lobe compartments, each component thus corresponding to only one or two cells per hemisphere [24].

**Fig. 6:**
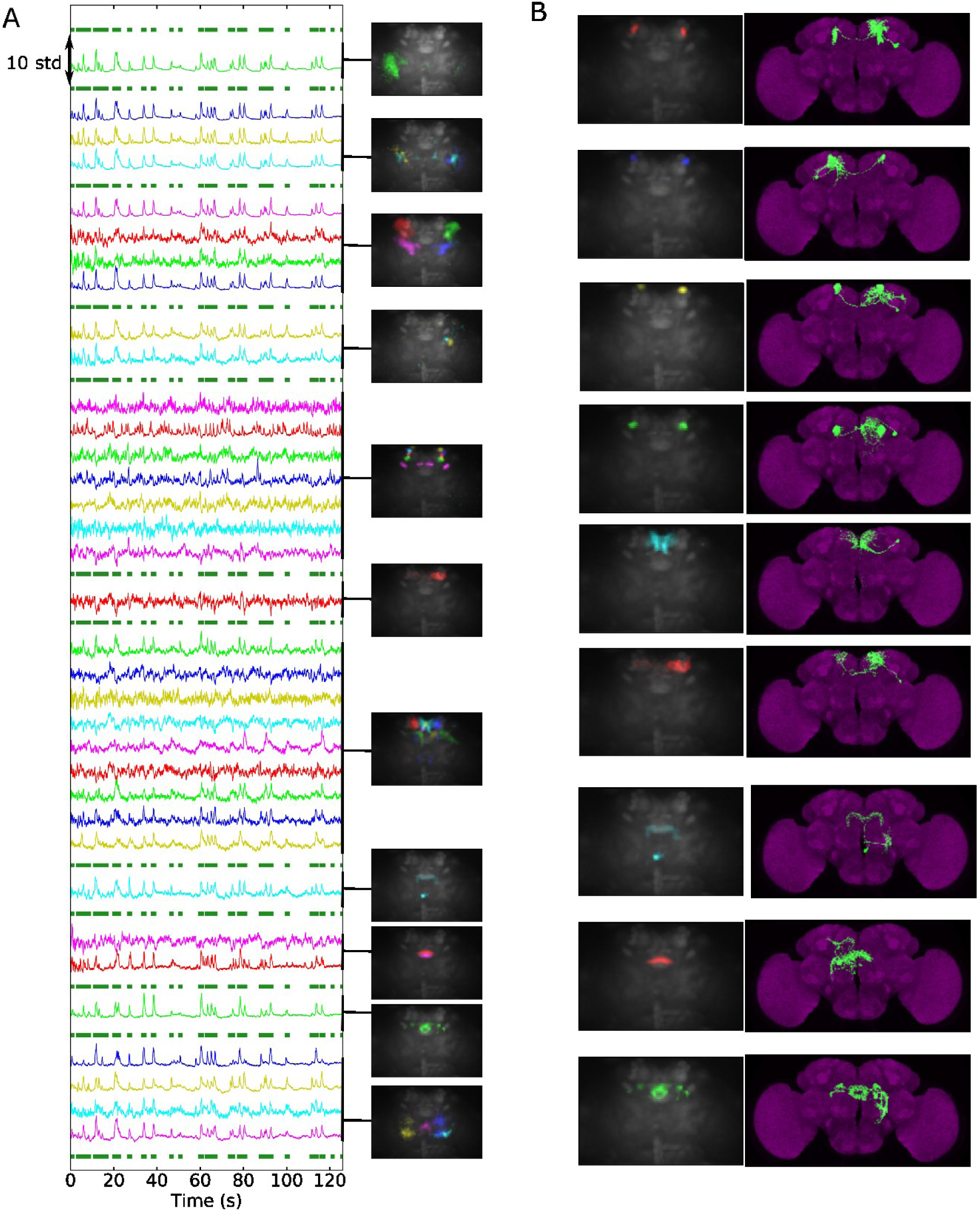
Components from flies expressing GCaMP6f in dopamine neurons (TH-Gal4 driver). A) All activity-related components are presented and sorted by brain region, with the color of the time series (which are variance-normalized) on the left, matching the color of the maps on the right (e.g., the first image corresponds to the first trace, the second image to the next three traces, and so on). Note that most components are strongly correlated with the fly walking (forest green traces interleaved with the component traces). The fly was resting or grooming the rest of the time. B) Example of TH-positive neuron from the Virtual Fly Brain database (right) matching the components’ maps (left). Note that the brain is tilted 20 degrees along the lateral axis compared to the template presented on the right. (See also Fig. S17).

### Components from pan-neuronal voltage recordings

As Figs. 7 and S19 demonstrate, voltage recordings with ArcLight also gave rise to maps portraying specific neuropils (and clearly distinguishable from artefacts as shown in Fig. S20). As Fig. S21 shows, the number of components per brain region was typically smaller than it was for GCaMP6–we extracted an average 174 (std=68, N=12) activity-related components (i.e., not noise or movement artefacts as detailed in the Methods section) from GCaMP6 recordings and 54 (std=14, N=6) from ArcLight recordings, probably because of the probe’s lower signal-to-noise ratio. However, ArcLight components were similar to those found with GCaMP6: in the optic lobe, some components responded to the onset and/or offset of light, with various degrees of adaptation. In the posterior slope, we found peaks at the onset of light. We also recorded large peaks of activity in the ventrolateral protocerebrum. Finally, we found components in the antennal lobe, lateral horn and mushroom body responded to odor (Fig. S22). The clearest difference between voltage and calcium was the presence in the ArcLight data of slow, spontaneous switches between the up and down levels of activity for components in a nodulus and contralateral protocerebral bridge (Fig 8). We did not observe those components in controls where GFP was expressed pan-neuronally. Furthermore, although some movements artefacts can generate slow fluctuations in those regions (see Fig 8B), we observed only the opposing or asynchronous switches for opposite sides of the brain when using ArcLight. Other patterns of spontaneous activity included slow oscillations in the antennal lobe and lateral horn, and fast, ongoing activity in the ellipsoid body rings and the protocerebral bridge (see Fig. S23 for an experiment in which many time scales of spontaneous activity were detected). More work is necessary to establish the conditions and consistency of these patterns of activity.

**Figure 7:**
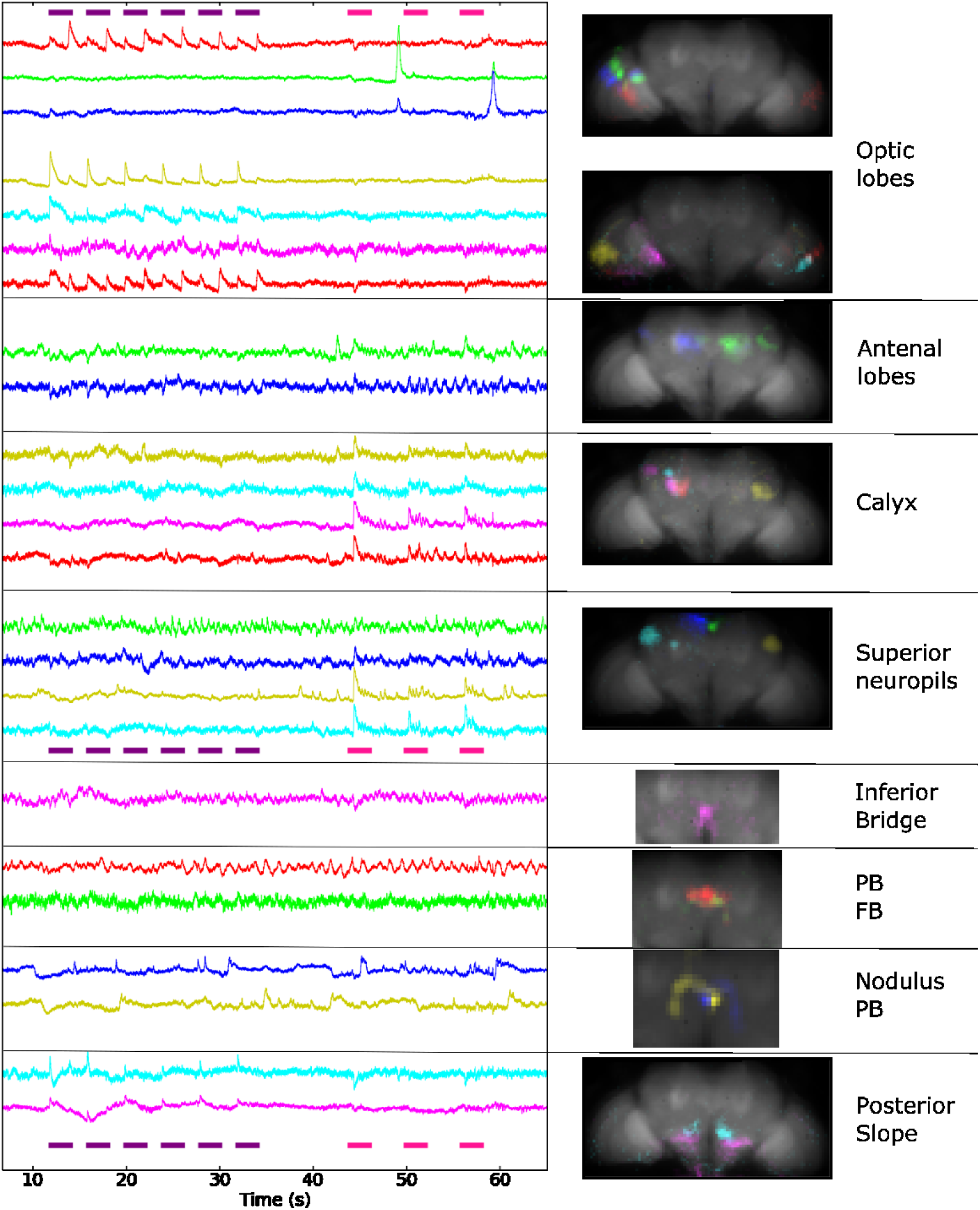
Components extracted from voltage activity. ArcLight was expressed pan-neuronally (nsyb-GAL4). The fly was presented with periodic flashes of UV light (violet bars) and puffs of apple cider vinegar (pink bars). The component time series are shown on the left (variance normalized), and the corresponding maps are on the right (PB: protocerebral bridge, FB: fan-shaped body), sorted by the brain region that was majorly present in the map. Note that the coronal plane was tilted 37 degrees away from the horizontal plane for this fly. See also Figs. S18 through S22.

**Figure 8:**
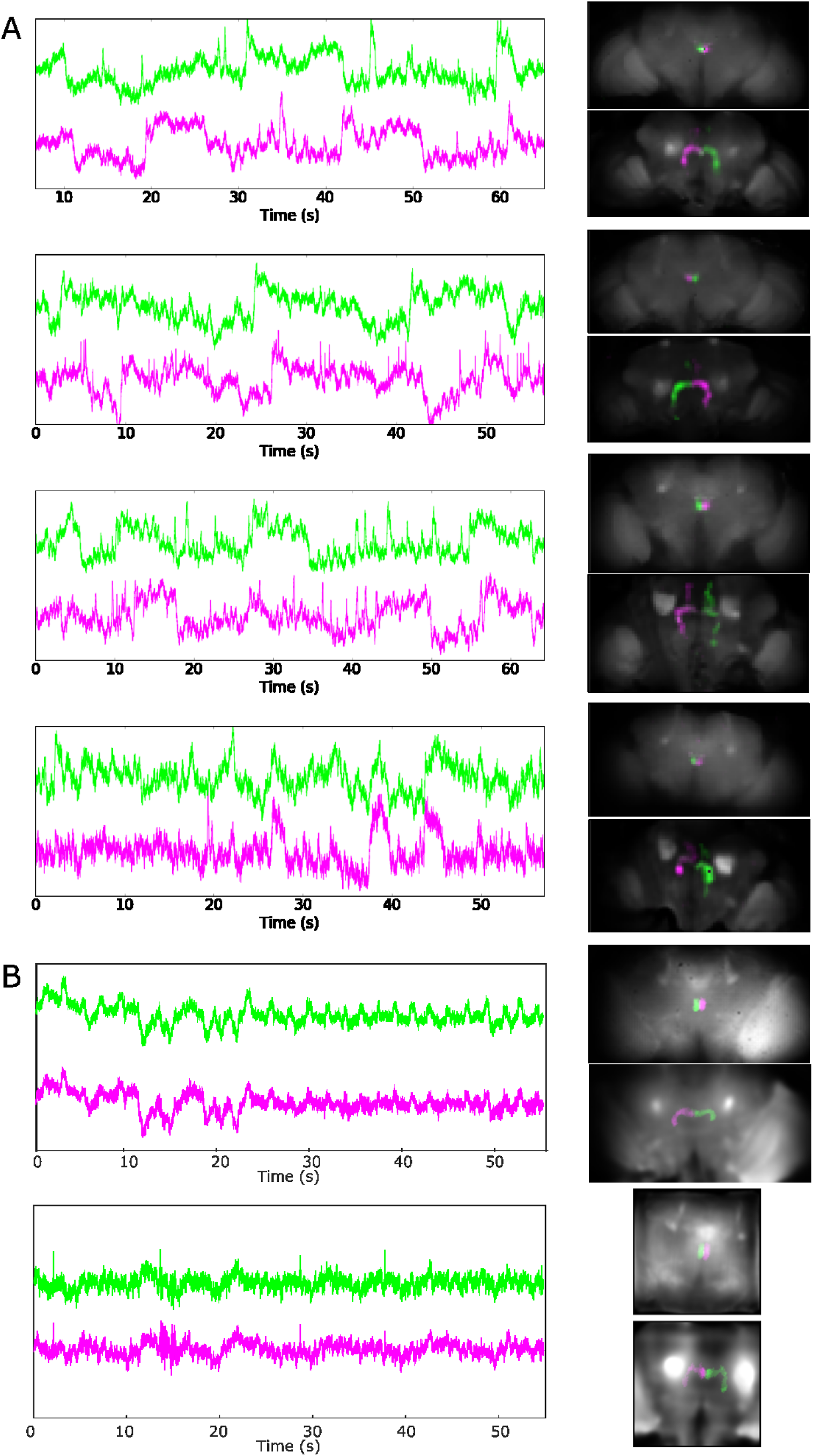
A) Spontaneous switches between up and down activity states for components in a nodulus and the contralateral part of the protocerebral bridge. We detected these components in all four flies studied (genotype: UAS_ArcLight, and from top to bottom: nsyb-Gal4, nsyb-Gal4, Cha-Gal4 and GMR57C10-Gal4, nsyb-Gal4). The images on the right present the two components at two different Z planes (at the level of the noduli and fan-shaped body, and at the level of the protocerebral bridge). B) Control in which GFP was expressed pan-neuronally. Because no similar components were automatically extracted, we created masks of one nodulus and the opposite side of the protocerebral bridge using an anatomical template.

Note that time series from single trials had high enough signal-to-noise ratios to detect graded potentials (e.g., components in the optic lobe in response to the onset and/or offset of flashes of light), and spike-like signals (e.g. spontaneous activity and odor response for components in the antennal lobe, mushroom body and lateral horn (Fig. 7 and Fig. S22)), which is consistent with previous literature [20] [19]. Spike-like signals were particularly clear in a more restricted driver for dopamine and serotonin neurons (i.e. TH-Gal4 and DDC-Gal4), as Fig S24 shows.

### Comparison of light field imaging to other techniques for adult Drosophila large-scale imaging

Recent studies have shown that large-scale brain imaging in flies is possible with other imaging methods. Mann et al. [25] used a high-speed, two-photon microscope to image the brain with higher resolution but at a slower rate (1Hz) in the absence of stimuli or behavior. We applied our analysis pipeline to these data to compare it to the results from the light field microscope (see Fig. S25). Out of the 23 activity-related components we obtained (compared to the average of 174 for light-field (see above), eight could be interpreted as cell bodies that could not be extracted at this resolution from the light-field data. Other components—components covering the antennal lobe, mushroom body, and lateral horn; components in the pars intercerebralis; and components in the antennal lobe glomerulus, calyx, and lateral horn (likely representing activity from antennal lobe projections neurons)—had similar spatial distribution as component from the light-field data.

Faster techniques have also been used to image large scale activity in flies: [26] used a Bessel beam to image 25% of the brain at 3.6Hz, and [27] used a light sheet approach (SCAPE, or swept confocally aligned planar excitation) to image a large portion of the brain at 10Hz. To test whether the fast frame rate of the light-field method enabled detecting signals otherwise undetectable with slower imaging methods, we subtracted the data smoothed over 100ms, thus revealing only the activity above 10 Hz. All GCaMP6f datasets (N=7) maintained activity related components (e.g., Fig. S26). When we did the same analysis with ArcLight data, four out of seven flies had at least one activity-related component. Note that more information from fast activity could be present in the data, but the low signal-to-noise ratio makes it difficult to detect it with PCA/ICA.

## Discussion

In this article, we presented a method to record fast near-whole brain activity in behaving flies, offered unique datasets that the method produced, and provided two examples of analysis pipelines that highlight this technique’s advantages.

First, we showed that the technique helps characterize global state changes related to response to stimuli and behavior. In particular, when looking at global activity in response to stimuli from different modalities, we obtained activity increases in brain regions known to be involved in processing these stimuli. Furthermore, we found a global pattern of activation when the fly walked in comparison to when it rested. In contrast, the activation was highly localized (in the areas of the AMMC, wedge, and saddle) when it groomed. This is consistent with a recent large scale optogenetic study that reported that the manipulation of many regions throughout the brain altered walking, but only a small region— the saddle—altered grooming [28]. The global activation during walking could be mediated by neuromodulators, and we indeed found that most dopamine neurons were silent during resting or grooming, but that many of these neurons, distributed over the whole brain, were active during walking. Dopamine activity could affect processing of underlying circuits (e.g., changing gain[29]), but could also be important for operant learning.

Second, we used a blind source separation algorithm to extract spatially distinct sources of activity. These sources matched anatomical sub-neuropil regions and thus helped identify neuron classes and sometimes a specific neuron type involved in the processing of specific stimuli or behavior, but also detected areas that were spontaneously active in the absence of changing stimuli and behavior. For example, we extracted activity in a nodulus and opposite side of the protocerebral bridge that switched between up and down states of activity. Although the fly was not walking in some of these experiments, the maps were similar to the pattern of expression of genetic drivers known to be involved in the fly’s handedness [30]. The time series resembled the flip flop pattern that was measured in the moth with intracellular recordings from neurons in the lateral accessory lobe [31], which is directly connected with the protocerebral bridge [15]. Note that the ability to record membrane voltage signals in the actual neuronal processes [32–34] performing the computation is an advantage over patch clamp experiments, which can only be recorded from the cell body at the periphery of the brain in Drosophila and thus may not be representative of the activity in the neuronal processes [35].

The datasets (available on CNCRS.org) of near-whole brain imaging and behavior contain additional, unexplored patterns of activity. Other analysis techniques will be necessary to extract all meaningful information.

The current method has several limitations that would benefit from being addressed in future work.

### Temporal resolution

The method permits imaging of the near-whole brain with a frame rate of 200 Hz; however, the time response of the probes we used in this study is slower (rise time to spike peak of ∼0.15 seconds for GCaMP6f and ∼0.1 seconds for ArcLight in our hands). Fast activity (i.e., at the onset of stimulus response or components with activity above 10Hz in Fig S24) can still be detected, but the probe’s response imposes a temporal filter on the underlying activity. The ability to record high signal-to-noise transients with such a high frame rate suggests that the light field microscope will be suited to measure activity from faster probes. This will help bridge the gap between the current fast local methods using microelectrodes (e.g., recording spikes and fast oscillations), and slower large-scale methods, such as calcium imaging.

### Effect of the excitation light

The excitation light excites the eye photoreceptors, thus affecting the fly’s ability to see, as well as potentially changing brain activity states. In the future, the fly’s blue light receptors could be genetically removed and potentially be replaced by another receptor, such as one for UV light, if a future researcher wanted to study brain responses to visual inputs without the strong background activation from the excitation light. For applications that do not necessitate visual inputs, blind fly mutants (e.g., a norpA; cryptochrome mutant) could be used to affect all light detection in the brain. Finally, one of the recently developed red probes could be used instead of GFP-based probes. Note that besides activation at the onset of excitation light, we observed two types of artifacts when the eyes were not completely protected from the excitation light. First, we observed sudden discharges in medulla column projections to the lobulla layers (see Fig. 7, second and third traces). Second, we observed oscillating waves propagating onto the medulla and along the lobulla in some calcium recordings instances.

### Effect of the preparation on the state of the fly

Dissection could have affected the fly’s state. Removing the cuticle on the back of the brain could affect brain activity by activating nociceptor neurons (e.g., those in the bristles). Dissection could also have affected the fly’s global health state. Indeed, we found that flies expressing ArcLight pan-neuronally were usually less active after dissection than they were before. Finding the optimal recovery time after dissection could help minimize these effects. Imaging non-dissected flies genetically modified to have a cuticle with low absorbance (e.g. yellow flies) could also help characterize the effects of the dissection.

Although the fly could still move its legs, abdomen, and (to a limited extent) its wings, the immobilization of its head, proboscis, and thorax could have affected brain activity and behavior by imposing unnatural constraints. Furthermore, the fly’s head was tilted more than 45 degrees in comparison to its natural position, in order to better align the thinner part of the brain to the z-axis. This helped minimize the loss of resolution with depth. We observed a seemingly natural behavior in this configuration (with alternations between grooming and walking as free flies do); however, we sometimes found the fly displaying unnatural behaviors, such as pushing the ball away or touching the holder with its legs.

Another problem resulting from immobilizing the fly’s head was the lack of coupling between the stimuli position and the fly’s movement that would normally occur in a natural setting. This problem can be solved using a virtual reality set-up in a closed loop configuration (e.g., using the movements of the ball for example to change the stimuli position).

### Data production rate

The whole procedure made it impractical to obtain data from a large number of flies. Even with practice, fly preparation remained challenging to the extent that it was difficult to obtain more than one good preparation per day. Another factor limiting data production was the reconstruction step, which takes approximately 10h on a cluster of 16 GPUs for a dataset of 60GB (which corresponds to approximately 1 minute of recording at 200Hz). This method is thus suited for studying complex spatiotemporal patterns and identifying neurons and brain structures in a few trials and flies, but not for larger studies such as genetic screens. Detecting sources from the raw light field data could help reduce the cost of reconstruction. For example, anatomical maps could be transformed back to a light field image and used as seeds for the source extraction algorithm [8].

### Detection of neuropil regions and neurons

Although we can observe the whole central brain (though the access to the gnathal ganglia depending on the quality of the preparation), as well as a large part of the optic lobes (typically the lobula and most of the medulla), we cannot observe all of the fly neurons. In particular, the ventral cord in the fly thorax is not accessible with the current set up. Imaging the ventral nerve cord might be feasible with the appropriate dissection preparation and an objective with a larger field of view.

As the brain contains approximately ∼10^5^ neurons, and we record, at most, a few hundreds of activity-related components, we are far from obtaining recordings from all neurons. This could be due to various reasons. First, some neurons might be silent and thus undetectable by our algorithm, which is based on temporal changes. Second, a number of neurons might contribute to one component. Indeed, the resolution of the microscope is, in general, larger than individual somata and neurites are, and neurites with similar presynaptic inputs and thus similar activity patterns will likely have similar geometry, making them indistinguishable to the algorithm. Additionally, the low axial resolution far from the focal plane makes it difficult to sort out the activity from regions that are close to each other such as the antennal lobe and the lateral accessory lobe, or the protocerebral bridge and the antler. However close to the focal plane, the functional maps were the same scale as functional maps obtained with higher resolution microscopy techniques were (e.g., in the fan-shaped body [36] and the lateral horn [37], respectively), or regions known to be functional units (e.g., antennal lobe glomeruli and ellipsoid body wedges and tiles). This suggests that the light field resolution might not limit the detection of functional units in comparison to higher resolution techniques, at least close to the focal plane. Third, neurons can be connected via gap junctions, making their activity too similar to be separated into different components. Using a second color and complementary drivers (e.g., drivers for excitatory versus inhibitory neurons, or drivers for main neurotransmitters versus drivers for neuromodulatory neurons) could increase the number of components that can be detected. Fourth, the signal-to-noise ratio might be insufficient for PCA and ICA to detect the activity in some processes. To increase the signal-to-noise ratio and obtain more components, future researchers could use a more sensitive probe (albeit at lower temporal resolution with GCaMP6s for example), record longer time series, or use faster probes to obtain more temporal information. Finally, the dimensionality reduction carried out with PCA might result in a loss of information that could be captured using other dimensionality reduction techniques.

The identification of anatomical structures could also be improved. Currently, the registration of the maps with the anatomy is done using landmark registration. This method is imprecise in brain areas that lack clear landmarks such as the ventral areas. Concurrently imaging the brains using a different microscopy technique with higher resolution could help detect more landmarks or make it possible to use different registration techniques. Automating the search for matches between the maps and neurons in large databases such as Flycircuit or Virtual Fly Brain would help to get to the level of neurons rather than brain regions.

The maps obtained using PCA and ICA can have regions with both positive and negative values, but this study has ignored the negative parts of the maps. More work is necessary to characterize the meaning of the negative values of the maps. In particular, neurons underlying the positive part of the maps could be inhibiting the neurons underlying the negative part of the map.

### Time series interpretation

The PCA/ICA algorithm used here helps to unmix neural activity from movement artefacts, or from other overlapping but different processes as well as scattered activity coming from other parts of the brain (see Figs. S11, S12, S14 to S16, and S20). However, the interpretation of these time series is not straightforward as there is no guarantee that the algorithm will extract the full neural activity from one region. Furthermore, the imperfect spatial separation of the sources can lead to artefacts when strong synchronous fluorescence changes are present in large parts of the brain. For example, in Figure S14B, a negative signal is present for a component in the optic lobe after the odor is presented. As this decrease is not present when measuring fluorescence in the region of interest delimited by the z-scored maps, or when applying PCA and ICA in the region of the optic lobes only, it is likely due to an imperfect separation of the optic lobe components from the regions in the middle of the brain where fluorescence strongly changes in response to the odor. Indeed, the maps for the optic lobe components have small negative values in the mushroom body and antennal lobe areas. To recognize these artefacts, observing both the unmixed time series and the region of interest fluorescence is thus advisable, as done in Figs. S12 and S14 to S16. Using a different algorithm such as non-negative matrix factorization might help prevent these artefacts, however, in our hands, the components obtained with non-negative matrix factorization were less localized, and thus more difficult to interpret than with PCA/ICA.

Movement correction with 3dvolreg can be imperfect and even can, in some cases, introduce additional artefacts when strong fluorescence changes are present in large parts of the brain. Furthermore, the algorithm uses rigid registration and does not correct for local deformations. Although we partly these artefacts at the PCA and ICA stages of the analysis, they can complicate the interpretation of some of the time series. Better movement correction methods with a limited sensitivity to fluorescence changes (such as RASL [7,38]) and non-rigid registration [39–41], as well as using an activity independent fluorophore in another color channel as a reference, would improve the reliability of the time series.

### Conclusion

Despite these limitations, the methods presented in this study can be used as a functional screen to identify brain regions and neurons involved in processing any stimulus or behavior that a fly can perform under the microscope. Furthermore, complementary to screens using activation or silencing of specific neurons, the voxels, regions and component’s time series give insight into the dynamics of the network, including ongoing spontaneous activity. This will help us understand how the brain accomplishes various functions, in particular those involving recurrent loops, such as integrating stimuli with various types of memory to guide behavior [24] and situating the animal in space [15,42].

## Methods

### Fly rearing and preparation for imaging

The fly genotype was as described in the figure legends, and fly stocks were obtained from the Drosophila Bloomington Stock Center, Bloomington, IN. Flies were reared at 25 ^o^C with a 24 h light/dark cycle on brown food (containing cornmeal, molasses, yeast, soy flour, agar, proprionate and nipogen), which has lower auto-fluorescence than yellow food (such as the one from the Bloomington Stock Center which contains yellow cornmeal, malt extract, corn syrup, yeast, soy flour, agar and proprionic acid).

Fly holders were 3D printed using Supplementary data chamber.stl. A piece of tape (Scotch 3M 0.75” wide) was shaped as a 1 mm high step using a 1 mm thick glass slide, and an aperture as is shown in Supplementary Figure 1 (1mm wide for the body and 0.6 mm wide for the head) was made by hand using a sharpened scalpel or a thin lancet (36 gauge, from TiniBoy). The tape was then stuck onto the chamber, aligning the opening of the tape to the center of the holder. We added nail polish at the contact between the tape and the holder to avoid leaks. We also added black nail polish (“Black Onyx” nail laquer, OPI products) to the tape to block the excitation light from hitting the fly’s eyes.

Note that although the black painted tape protected the flies’ eyes from direct illumination by the microscope’s excitation light, the light scattered by the brain can still activate the eye’s receptors for blue light, as the transient activity in the first few seconds of each experiment demonstrates (see for example the optic lobe trace in Fig. S6). To verify that these receptors were not saturated we presented flashes of 470 nm blue light as external stimuli (See Fig. S27). Although the stimuli excited fluorophores non-specifically, PCA and ICA could still extract neuronal calcium responses in the optic lobes and the protocerebral bridge, thus demonstrating that the fly could still perceive external blue stimuli.

At the start of an experiment, flies were transferred to an empty glass vial and left on ice for approximately one minute. The holder was put in contact with wet tissues on ice under a stereomicroscope. A fly from the cold vial was pushed into the holder’s opening so that the posterior part of the head was in contact with the tape. UV-curing glue (Fotoplast gel, from Dreve), was added at the junction between the tape and the head between the eyes and cured for 5 seconds using a 365 nm Thorlabs LED light at 20% of power for 5 seconds. A piece of thin copper wire (wire-magnet, 40 gauge, from AmplifiedParts) or a piece of tape was placed above the legs to push them away from the proboscis (see Fig. S1). UV glue was then added at the rim of the eye and all around the proboscis (which was pushed into the head), without touching the antenna or the legs, and was cured for 5 seconds. Uncured glue was carefully removed with tissues. A small amount of vacuum grease was placed around the neck behind the proboscis to avoid later leaks. The wire or tape was then removed, and a small piece of tissue paper or a small Styrofoam ball was given to the fly to walk on to monitor its health during the following steps.

The holder was turned over and the fly’s thorax was pushed down to clear the way to the back of the brain. Small pieces of tape were added onto any remaining holes around the fly’s body, and UV glue was added on top of them and cured around the thorax to fix it in place. Vacuum grease was then pushed toward the neck with a tissue. Saline (108 mM NaCl, 5 mM KCl, 3 mM CaCl2,4 mM MgCl2, 4 mM NaHCO3, 1 mM NaH2PO4, 5 mM trehalose, 10mM sucrose, 5 mM HEPES adjusted to pH 3.35 +/−0.05 with NaOH, prepared weekly) was added and left for few minutes to make sure that there were no leaks.

Fresh saline was added, and dissection was started with forceps (#5SF, from Dumont) that had been previously sharpened as finely as possible by hand. We first removed the cuticle in the middle of the back of the head, being careful to cut pieces before pulling them out. This exposed the hole in the middle of the brain where muscle 16 resides. The pulsatile piece was pulled out. Fresh saline was added, and the remainder of the cuticle was removed piece by piece. The brain was washed with saline several times to remove fat bodies. The air sacs were then removed very carefully as to try not to displace the brain. After a new wash with saline, the fly was ready for imaging.

### Imaging set up

The microscope was modified from an upright Olympus BX51W with a 20x NA 1.0 XLUMPlanFL, or a 40x 0.8 NA LUMPLFLN objective (from Olympus). A microlens array with pitch=125μm and f/10 to match the 20x objective, or f/25 to match the 40x objective [3] (from RPC Photonics), was positioned at the image plane using a custom made holder (with some parts from Bioimaging Solutions, Inc). Two relay lenses (50mm f/1.4 NIKKOR-S Auto, from Nikon) projected the image onto the sensor of a scientific CMOS camera (Hamamatsu ORCA-Flash 4.0). Note that when using half the camera frame to attain 200Hz for voltage recordings, the brain fit within in the field of view, but rays coming from points far from the focal plane with a large angle were missed, slightly impairing reconstruction. A 490nm LED (pE100 CoolLED) at approximately 10% of its full power was used for excitation. We used a 482/25 bandpass filter, a 495 nm dichroic beam splitter, and a 520/35 bandpass emission filter (BrightLine, Semrock) for the fluorescence. We measured the power at the sample with a power meter and found that it was up to 1mW for the 40x objective and 4mW for the 20x objective. Photobleaching led to a decrease in intensity after 30 seconds of 13% (N=12, SD=9%) for GCaMP6, and 20% (N=6, SD=13%) for ArcLight. Note that the full set up cost approximately USD $37000 (USD $52000 with the 64Gb of RAM acquisition computer and the 256Gb of RAM analysis computers), which was substantially cheaper than other cutting-edge microscopy techniques such as two-photon microscopes are.

The resolution as a function of depth (see Fig. S2) was determined by imaging 2 μm fluorescent beads dispersed in an agarose gel. After reconstruction, the center of beads at different distances from the focal plane were recorded using ImageJ, and a MATLAB program measured the point spread function’s axial and lateral full-width at half-maximum (see https://github.com/sophie63/FlyLFM for the code).

The lateral field of view for the 20x objective was 624 × 636 square microns (312 × 309 for the 40x objective), as was determined using a mire.

The fly holder was positioned on a U-shaped stage above an air-supported ball so that the fly could walk (see Fig. 1 and supplementary videos). The ball was either polyurethane foam (10 mm in diameter), Styrofoam, or hollow HDPE (1/4 inch). We prepared a cup matching the ball diameter and with a 1.2 mm hole using self-curing rubber (from Sugru). A bottle of compressed air provided a steady flow in a pipeline consisting of a tube and a pipette tip connected to the cup hole. A micromanipulator (from Narishige) positioned the ball under the fly’s legs. For some flies, we instead provided a small Styrofoam ball that the fly could hold. The fly and the ball were illuminated by a row of IR LEDs (940nm, from RaspberryPiCafe®) in front of the fly, and were observed at 100 Hz using a small camera (FFMV-03M2M, from Point Grey).

To better align the behavior with the fluorescence, in some experiments the camera for monitoring the behavior and the fluorescence were synchronized by using the output of the Flash4.0 camera to trigger the acquisition from the behavior camera. When imaging the fluorescence at 200Hz, one triggering signal out of two was ignored by the slower behavior camera that recorded at 100Hz. We recorded the fluorescence images were recorded with HCImage (from Hamamatsu) and streamed them to RAM on a 64Gb of RAM computer, which allowed us to record approximately one continuous minute.

For the odor stimulus, air was delivered by a pump (ActiveAQUA AAPA7.8L) through an electrically controlled valve (12 Vdc normally closed solenoid valve, from American Science & Surplus), bubbled in 50% ethanol or 50% apple cider vinegar in a vial, and blown towards the fly through an inverted 1 mL pipette tip. The valve circuit was controlled by a relay (RELAY TELECOM DPDT 2A 4.5V, Digi-Key), that was connected to a LabJack U3-HV through a LJTick-RelayDriver (from LabJack). For visual stimulation, the excitation light and a 365nm or 470nm LED (from Thorlabs), were also triggered by the LabJack, which was controlled using MATLAB programs (see https://github.com/sophie63/FlyLFM for the code).

### Analysis

We reconstructed the light field images were reconstructed using a program in Python as described in ref. [3]. Briefly, a point spread function library corresponding to the specific set up was first generated: we typically chose to reconstruct a stack of 40 layers (separated by 6 microns), with a lateral sampling distance of either 3 or 6 microns. The voltage probe’s low signal-to-noise ratio made it more difficult to detect signals with a finer sampling, so we typically reconstructed the voltage data with a lateral sampling distance of 6 microns and the calcium data with a lateral sampling of 3 microns. We reconstructed the images using 3D deconvolution on a cluster of GPUs (generously provided by the Qualcomm Institute at UCSD). Note that reconstruction using cloud computing (AWS) would cost ∼$0.003 dollars per volume. A dataset of 10000 time steps required approximately eight hours to reconstruct on a cluster of 15 GPUs.

We assembled the images in a Nifti file using a python routine (Tiff2niiV2 in https://github.com/sophie63/FlyLFM), inspected and cropped them in FIJI, and discarded the first 5 seconds because the strong activity in response to the excitation light made it difficult to correct for movement and photobleaching. The 3D image registration function 3dvolreg [43] from AFNI was then used to correct for rigid movements. We removed the background fluorescence and the decrease in intensity from photobleaching by subtracting a signal smoothed using a box average over 15 to 30 seconds depending on the severity of the bleaching and the length of the recording. The time series were then multiplied by −1 for ArcLight data. For denoising, we found that a Kalman filter (from https://www.mathworks.com/matlabcentral/fileexchange/26334-kalman-filter-for-noisy-movies) with a gain of 0.5 was better than a median filter over 3 points was, and we used this for the figures in this paper. We then applied SVD to subtract components that were most clearly related to movement: their maps contained shadows around areas with different background intensities as shown in Fig S4. For some flies, different conditions corresponded to different datasets, which we concatenated in time after preprocessing. The reconstructed data as well as the data after preprocessing will be soon available on the CRCNS website (https://crcns.org/NWB/Data_sets).

For early datasets (before direct synchronization of the cameras), the fluorescence and the behavior were aligned using the onset and offset of the excitation light. The small discrepancy (approximately 30 ms per minute) between the total time given by the camera for the fluorescence and the camera for the behavior was corrected linearly, then the fluorescence data was interpolated to match the behavior data using the MATLAB function Interpolate2vid (in the cases for which the cameras were not synchronized).

We manually analyzed the behavior (noting the times of the behaviors, or pressing different keyboard keys when we recognized different behavior using the MATLAB GUI Video_Annotate in https://github.com/sophie63/FlyLFM). We also characterized the fly’s walk by tracking the movements of the ball using ImageJ’s optic flow plugin (Gaussian Window MSE).

Maps comparing the activity during rest and walking, resting and grooming, turning left and turning right, and one second after stimulus presentation compared to one second before were obtained by simply averaging the time series in each voxel for the different conditions, and subtracting these maps from one another. The positive value was colored in magenta and the negative in green thus showing which condition dominated in which voxel.

The average of the volume time series was aligned to an anatomical template (available here: https://github.com/VirtualFlyBrain/DrosAdultBRAINdomains) in which the brain is segmented into regions according to the nomenclature described in ref. [14]. The registration was performed using landmarks with ImageJ (as described in http://imagej.net/Name_Landmarks_and_Register). We marked several points in the protocerebral bridge, the tips of the mushroom body (between the alpha and alpha’ lobes), the middle of the noduli, the lateral tip of the lateral triangles, the lateral tip of the lateral horns, the center of the ellipsoid boy, the center of the antennal lobes, and the bottom part of the trachea at the level of the noduli. Although the landmarks were readily observable with the background fluorescence (see Fig S3 for example), making a template superposing the components to the volume average helped to visually find the landmarks.

Dimensionality was then reduced by separating the volumes into slices of thickness corresponding to the point spread function height, and averaging in z for each slice. The 4D datasets were typically (x,y,z,t)=200×100×10×10000 at this stage.

For source extraction, we found that using melodic [12] from the FSL package directly gave meaningful components. However as the code was running slowly on our large datasets, so we adapted it in MATLAB to parallelize some steps. A first step of SVD was used to remove the largest part of the components before calculating the variance that was used to normalize the data. We then reapplied SVD and plotted the log of the singular value spectrum to automatically detect the shoulder at the point with a 45° tangent. We found that although the components with the smallest variance were noise, some activity-related components were still present after the shoulder point. As such, choosing twice the number of components at the shoulder gave a good compromise between keeping activity and removing noise components. ICA was then applied with FastICA [44] (see ICAalamelodic.m file from https://github.com/sophie63/FlyLFM). The sign was chosen so that the average of the positive side of the map was larger than the negative side. The components were then automatically sorted by brain region: we averaged the components maps in anatomically defined regions[14] using regions masks, and chose the main region as the one with the highest average. We removed components corresponding to movement or noise partly automatically (removing components present in more than 5 regions and containing more than 200 separate objects) and partly by hand (see example of typical artifactual components in Fig. S11) using a Jupyter notebook (the notebooks corresponding to the choices made for the figures in this paper can be found at https://github.com/sophie63/FlyLFM).

To obtain the region of interest time series, we first made masks using the PCA/ICA maps. We calculated the standard deviation from the value in the map, and set all voxels with values inferior to 3 times that std to zero. We then used those mask to do a weighted average of the ΔF/F time series.

The time series for turning left or right were obtained by convolving the optic flow for the ball going left or right, with a kernel corresponding to the impulse response of GCaMP6. These time series were then regressed with the components’ time series, and we inspected the maps with the strongest regression coefficients.

### Image manipulations

The Fig. 1B bar was added with ImageJ and the 3D rendering was done in Icy [45], in which transparency and contrast were adjusted globally on the volume. The component’s maps were thresholded at 3 × σ, and only the positive part of the map was displayed. The image contrast was then globally adjusted in ImageJ, and the figures panels were assembled in Inkscape.

### Code availability

The MATLAB and Python code for preprocessing, PCA/ICA and sorting of the components is available at https://github.com/sophie63/FlyLFM.

## Author Contributions

R. Greenspan and T. Katsuki had the idea of using light field microscopy to image the whole fly brain. T. Sejnowski proposed to use PCA and ICA to extract activity from the recordings. T. Jia performed data analysis and curation. L. Grosenick, M. Broxton and K. Deisseroth provided advice and training on the light field microscopy setup, the code for 3D deconvolution, and a computational infrastructure for deconvolving large numbers of volumes. R. Greenspan and T. Sejnowski supervised the research. T. Katsuki performed early experiments and advised S. Aimon throughout the project. S. Aimon designed and performed the experiments presented in this paper, did most of the analysis, and wrote the manuscript. All the authors critically reviewed the manuscript.

## Acknowledgements

We thank Jürgen Schulze, Joseph Keefe and Calit2 for generously providing a GPU cluster. Charles F. Stevens contributed many helpful discussions and comments on this manuscript. Angelique Paulk provided valuable advice on the floating ball setup. Stephen M. Smith helped with reproducing parts of melodic in MATLAB, Ryan Shultzaberger helped with R and bash, Daniel Soudry helped with source extraction, and Nicolas Aimon helped with fitting time series in python. Alexander Pope helped with fly preparation. M. Nitabach and V. Pieribone gave us UAS-ArcLight flies and the Nitabach laboratory provided technical support on imaging ArcLight.

